# Oscillatory multiplexing indexes precision

**DOI:** 10.1101/205245

**Authors:** Laetitia Grabot, Tadeusz W. Kononowicz, Tom Dupré la Tour, Alexandre Gramfort, Valérie Doyère, Virginie van Wassenhove

## Abstract

Oscillatory coupling has been implicated in the representation and in the processing of information in the brain. Specific hypotheses suggest that oscillatory coupling may be relevant for the temporal coding of information but to which extent this may translate to conscious timing is unknown. Here, we tested the hypothesis that the temporal precision of self-generated timed actions may be controlled by phase-amplitude coupling (PAC). Using a timing task, we show the existence of significant alpha-beta (α-β) PAC, robust at the individual level, and specific to temporal production. Second, an increase in the strength of α-β PAC was associated with a smaller variance in time production, i.e. an increased precision in timing, but there was no correlation with the duration of the produced interval. Our results suggest an active role for α-β coupling in maintaining the precision of the endogenous temporal goal during time production: specifically, α oscillations may maintain the content of current cognitive states, thus securing the endogenous temporal code for duration estimation instantiated in β band. Oscillatory multiplexing may thus index the variance of neuronal computations, which translates into the precision of behavioral performance.

## Introduction

The relevance of neural oscillations for cognitive operations has been acknowledged over the years (Buzsáki & Draguhn, 2004; Fries, 2015). Interactions across frequency regimes, referred to as oscillatory nesting or multiplexing (Canolty & Knight, 2010), likely supports long-range communication and integration over spatial and temporal scales (Akam & Kullman, 2014; Fries, 2015). A common form of observed oscillatory nesting is the modulation of high-frequency power (e.g. gamma, γ) by the phase of low-frequency oscillations (e.g. theta, θ). Such phase-amplitude coupling (PAC) has been well established in neuroscience (Tort et al., 2008; 2009) with mounting evidence that PAC is functionally relevant for the representation of temporal sequences (Heusser et al., 2016), working memory (Axmacher et al., 2010; Lisman & Jensen, 2013; Roux & Uhlhaas, 2013; Voytek et al., 2010) and speech processing (Canolty et al., 2006; Giraud & Poeppel, 2012). In particular, the observation that low-frequency neural oscillations regulate spike timing in the human brain has raised the hypothesis that PAC provides a temporal code for cognition (Jacobs et al., 2007; Buzsáki, 2010). Whether such temporal code impacts conscious timing is unknown.

Oscillatory multiplexing has been proposed to mediate the integration of information across temporal scales during working memory and interval timing (Gu et al, 2015). Although the precise manner with which oscillatory multiplexing contributes to time estimation remains unclear, a literal read would be that information content, encoded in higher frequency activity, would be integrated over the time scales of the low-frequency oscillations (van Wassenhove, 2016). This working hypothesis would effectively predict that the coupling strength in PAC should linearly increase with the length of the perceived or produced time interval. Such mechanism would be consistent with internal clock models (Treisman, 1963) in which perceived duration results from the *integration* of information (i.e. pulses or events count) over time. An alternative hypothesis would be that PAC regulates the *precision* of information during its transmission and/or maintenance over relevant brain regions. Specifically, it has been shown that temporal expectations can align the phase of endogenous low-frequency oscillations at the time at which sensory events were expected to occur (Samaha et al., 2015), reflecting a form of temporal optimization for sensory encoding. Θ-β PAC has been suggested by another study (Cravo et al., 2011) to be sensitive to the occurrence, or absence, of a predicted external event in time. Altogether, these studies suggest a link between temporal expectation of sensory events and PAC. However, whether such mechanism can be elicited endogenously (or volitionally) has not been addressed.

In this magnetoencephalography (MEG) study, we contrasted the *integration* and the *precision hypotheses* for PAC using an explicit time production task, and we questioned whether an increase in PAC would index timing performance *per se* (i.e. the duration of the produced interval), or timing precision. For this, we thus distinguished precision (i.e. variance of temporal production) from accuracy (i.e. variation of the temporal production relative to the target interval). We show that generating a time interval in the absence of sensory stimulation was accompanied by a robust alpha-beta (α-β) PAC observed at the individual level. Importantly, the strength of α-β coupling correlated with the precision and with the accuracy of timing performance, but not with the absolute produced duration itself. Our results thus provide support for the fundamental role of oscillatory multiplexing in temporal coding, extending this notion to conscious timing and performance precision.

## Results

### Behavioral evidence for variable precision and accuracy

12 participants were asked to produce 1.45 s and 1.56 s by briefly pressing a button at the start (R1) and at the end (R2) of their time production (TP). Participants complied with the task requirements by producing 1.513 s time intervals on the first three, and 1.614 s time intervals on the last three experimental blocks (Fig. 1B). Although the overall performance was quite accurate, the time production data showed large variability within-individuals (across blocks) but also across individuals (Fig. 1CD). This variability was quantified by the Coefficient of Variation (CV, Fig. 1C) and by the Error Ratio (ER, Fig. 1D): the CV was calculated by dividing the standard deviation by the mean duration production and the ER was calculated by dividing the mean time estimates by the target interval in a given block. Hence, we considered the CV as a measure of precision, and the ER as a measure of accuracy. Both metrics were calculated per experimental block and per participant. Although changes in feedback were manipulated in the experimental design, there was no *ad-hoc* hypothesis regarding its effects on possible PAC, notably because no significant changes in precision were found as a function of feedback as tested by repeated measures ANOVA (*F*(5, 65) < 1, *P* > 0.1). Additionally, as can be seen in Fig. 1D, a general trend for a lengthening in duration estimation was observed over the entire course of the recording: this drift was shown to not depend on changes in feedback or changes in duration (Kononowicz et al., *under review*). We then explored the role of oscillatory multiplexing in timing variability.

**Fig. 1.**
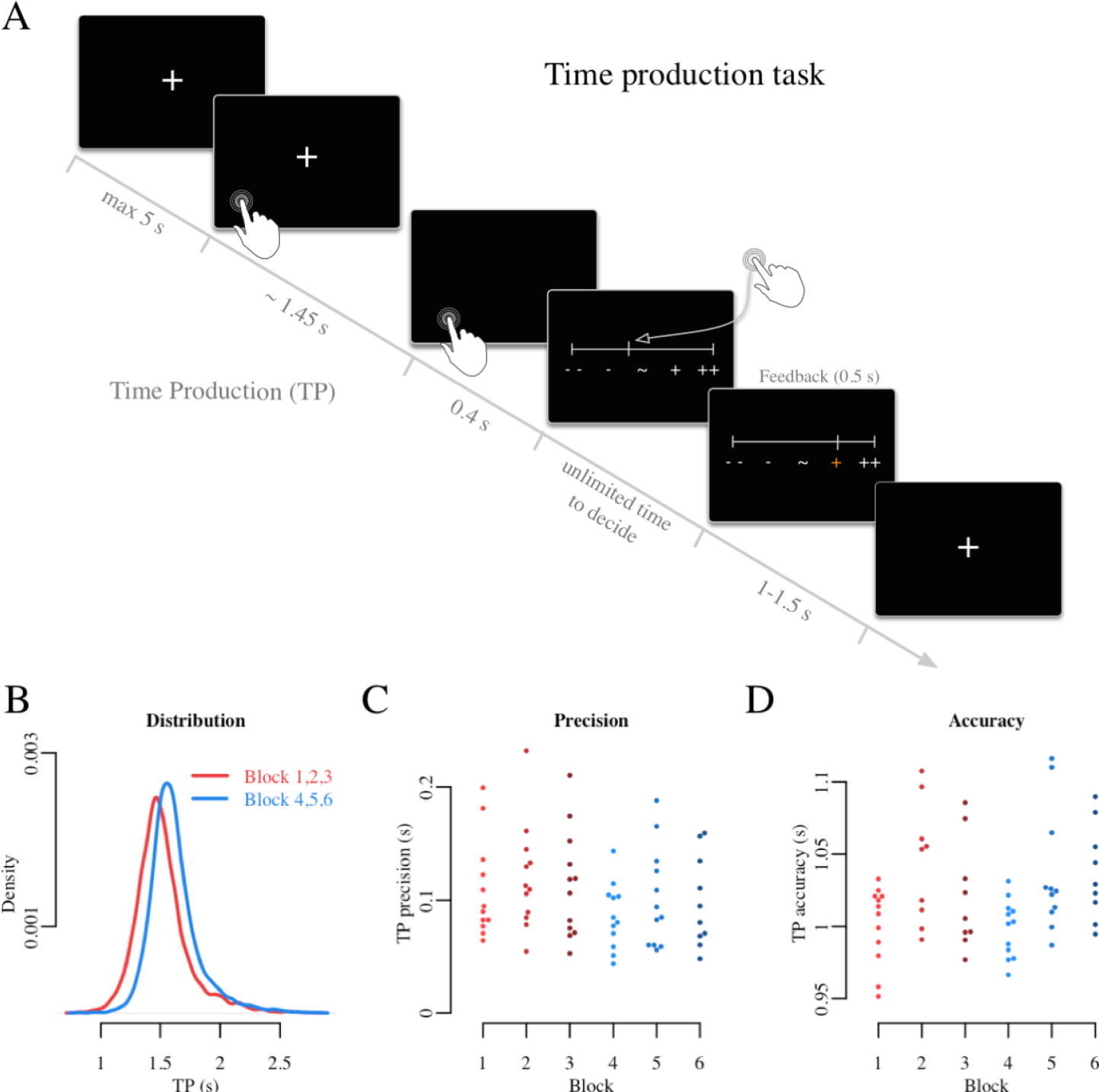
Probing behavioral precision and accuracy using a time production paradigm. (*A*) Time course of an experimental trial. (*B*) Probability density function of all time productions (TP) when producing 1.45 s (red) and 1.56 s (blue). (*C*) Temporal precision computed as Coefficient of Variation (CV) over time productions per block (color) and per individual (dot). (*D*) Accuracy in time production computed as Error Ratio (ER = *production/target*) per block (color) and per individual (dot).

### Robust α-β coupling during time production

Ahead of PAC calculation, we assessed the power spectrum during the timed interval (from 0.4 to 1.2 s following R1 to avoid evoked responses) by averaging the PSD across all sensors, conditions and individuals. Two peaks were readily visible in the spectrum at α (~10 Hz) and β (~25 Hz) frequencies, suggesting the existence of two plausible regimes for oscillatory multiplexing (Fig. S1A). The α PSD was mainly localized in occipital and parietal regions (Fig. S1B). To assess whether any form of PAC was present during interval production, we computed the modulation index (MI) in this same interval (Tort et al., 2009). Visual inspection of grand average data across all individuals and sensors revealed a strong MI between the α phase and the β power in centro-parietal sensors (Fig. 2B). For dimensionality reduction, we selected the 10 sensors for each individual displaying the maximal α-β MI: as can readily be seen in Fig. 2A, the 10 sensors with maximal α-β MI were found in central and parietal regions for a majority of participants. For each sensor, statistical significance of the MI was assessed at the individual level by shuffling the α phase (Tort et al., 2010). A Z-score above 4 (P < 0.0001; Fig. 2A for individual outlines), revealed that α-β PAC was statistically significant for each individual.

**Fig. 2.**
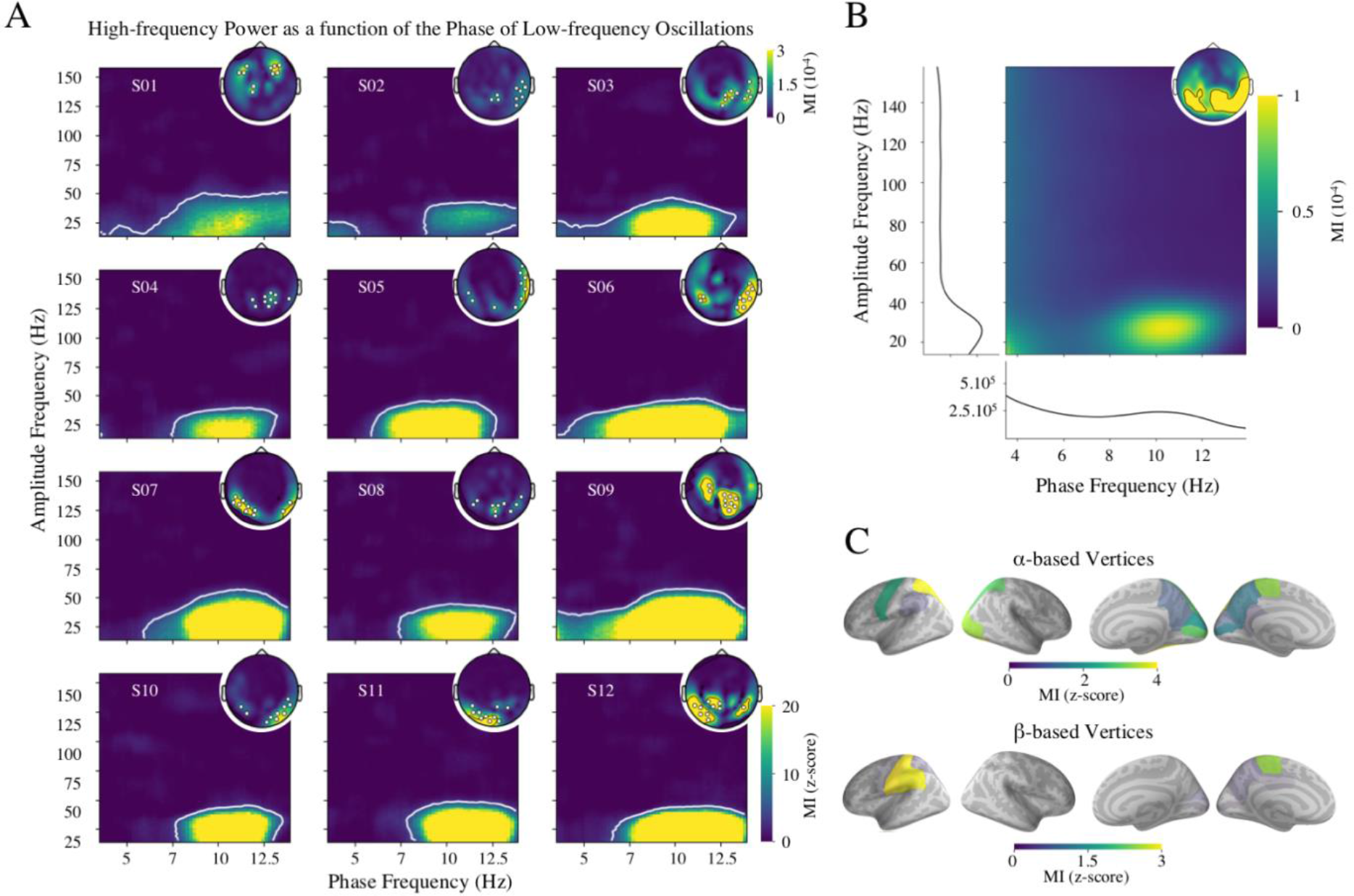
α-β PAC during time production. (*A*) The modulation index (MI) for each individual. A set of 10 sensors showing a maximal α-β MI was selected for each participant. The topographic plots (inset) map the α-β MI and highlight the selected sensors in white. The comodulograms show the MI for all analysed frequencies, averaged across selected sensors. The white outlines delineate significant Z-scored values above 4 (*P* < 0.0001). (*B*) The grand average comodulogram across all trials, participants and sensors, showing significant α-β PAC. The average topographic map of α-β MI plotted in the right inset shares the same scale as in panel A. (*C*) Z-score MIs were computed in cortex. Brain regions showing maximal MI on the basis of α or β power are highlighted in the top and bottom panels, respectively.

Across participants, α-β MI was maximal for the phase of the 10.3 Hz peak frequency (SD = 0.6) and for the amplitude of the 27.2 Hz peak frequency (SD = 3.9) (Table S1 provides all individual peaks). Both values were consistent with the frequency peaks observed in the previous PSD. Specifically, we observed that an individual’s α peak frequency also corresponded to the α peak frequency of the individual’s PAC [R = 0.74, P = 0.004; Fig. S3]. Additionally, the β peak in α-β PAC was distinct from the 2^nd^ harmonic of the α peak in the α-β PAC (t = 7.064, p < .001; CI_95_ = [4.5; 8.6]) meaning that the observed α and β peak frequencies were not simply harmonics.

To cross-validate our findings, and insure that the observed PAC was not spurious (Hyafil, 2015; Jensen et al., 2016), we quantified α-β PAC using a recently developed Driven Auto-Regressive (DAR) statistical modeling approach (Dupré La Tour et al., 2017) which makes no *a priori* assumptions on the shape of the signal. DAR modeling revealed a comparable α-β PAC (Fig. S2). Interestingly, this new method provided a narrower focus on higher frequencies of power modulation, suggesting slightly larger specificity of high power modulation. Noteworthy for both methods, the peak of power frequency (Tort method = 27.2 Hz; DAR method = 34.5 Hz, SD = 2.3) lied in the vicinity of β and lower γ frequencies, suggesting that for every α cycle at least one cycle of β was transiently modulated by the phase of α oscillation. This transient modulation may be related to the known ‘burstiness’ of beta oscillations, suggesting that on single trials, β oscillations may not be sustained but occur in a single cyclic burst of activation (Jones 2016; Sherman et al., 2016). As previously observed, the peak frequencies for α and β found in the α-β PAC with the DAR method showed no clear harmonic relationship (t = 18.641, p < .001, CI_95_ = [11.1; 14.0]).

To further assess which brain regions may exhibit the highest degree of coupling, we source-estimated PAC in cortex. First, we reconstructed the time-resolved signals on a singletrial basis, and used the same approach for PAC calculation as we did for the sensor data. We spatially defined brain regions from a known cortical parcellation using maximal *α* power as our criterion (see Material and Methods). α-β PAC was maximal in left sensorimotor regions, presumably due to the motoric components of the task (post-, pre-, para-central, and supramarginal areas) (Fig. 2C). High PAC was also found in parietal regions, in line with the notion that endogenous *α* rhythms would largely contribute to PAC. The α peak analysis (Fig. S1C) and the PAC source estimation (Fig. 2A) were thus topographically consistent with each other. To test the robustness of source estimations, we also conducted the analysis with a selection of spatial location based on the maximal *β* power: the left sensorimotor areas showed maximal α-β PAC, consistent with the *α* -based observations (Fig. 2C, bottom panel).

### α-β coupling is specific to the interval being timed

Although we showed that α-β coupling was present during the production of temporal intervals, one may argue that the observed coupling was strictly relevant to motor preparation as opposed to timing. To test this, we capitalized on our experimental design, which required that participants self-initiate their time productions. As participants volitionally initiated their first button press (R1) with no explicit time requirement or pressure, we used the data ranging from - 0.8 s to R1 as control for motor preparation. We specifically tested whether the MI during the production of the time interval was significantly increased compared to baseline (before R1). A cluster-based permutation t-test on comodulograms averaged across the selected sensors and across participants showed that α-β PAC was significantly larger during time production than during the volitional trial initiation (*P* = 0.037, Fig. 3). As the number of trials for PAC estimation was not optimal for 3 participants (Fig. S4A), the cluster permutation test was repeated for the 9 participants with a sufficient number of trials for robust estimation (> 40 trials) revealing again a PAC significantly larger than in baseline (*P* = 0.016, Fig. S4B).

**Fig. 3.**
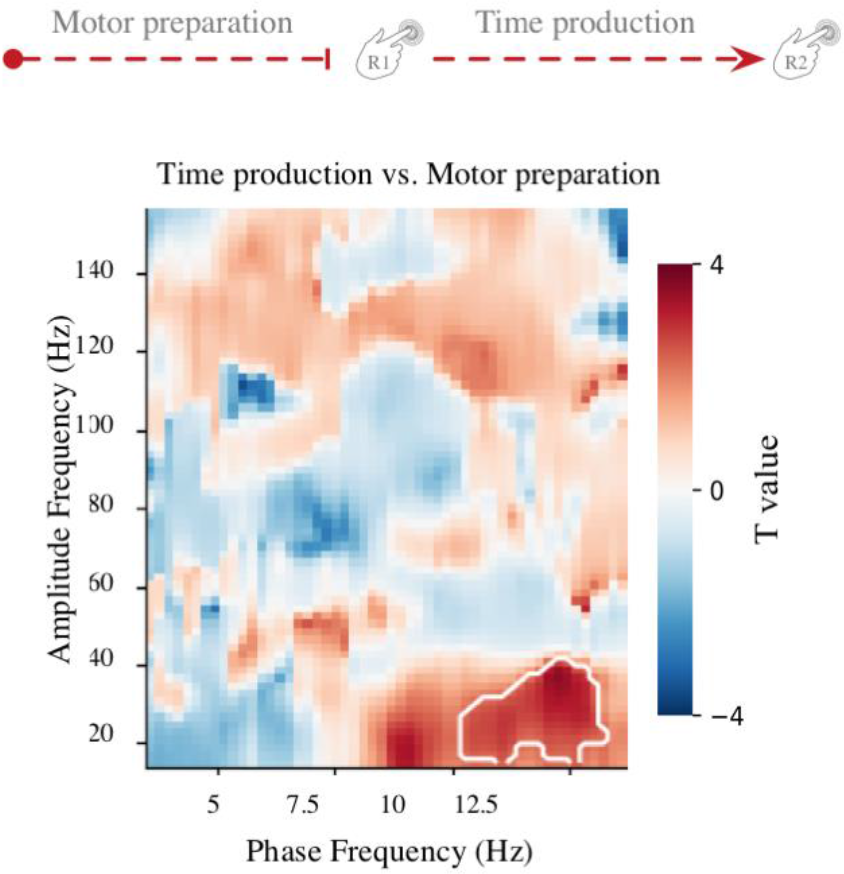
α-β PAC is specific to the timed interval R1-R2. To ensure that α-β coupling was related to timing processes, we contrasted PAC during the produced time interval (R1 to R2) with PAC during baseline of the motor preparation to the same trial (−0.8s to R1). α-β PAC was significantly higher during the time production interval as compared to motor preparation (corrected cluster delineated in white, P < 0.037).

Importantly, because the power of the low-frequency oscillation may impact the MI (Dupré La Tour et al, 2017), we also insured that *α* power density did not differ before the time interval and during the production of the time interval (*P* > 0.1). The α-β power ratio did not significantly differ between these two time periods (*P* > 0.1). In light of these controls, the α-β coupling could not be explained by a simple power difference between the two time periods. To put our findings in context, it is noteworthy that in a study requiring similar motor demands (De Hemptine et al., 2013), PAC was shown to decrease during motor preparation as compared to the movement phase; here, during a timing task, we found the opposite pattern.

Altogether, our comparisons against baseline showed that the observed α-β PAC were likely linked to the task requirements of endogenously producing a timed interval, thus could play a functional role in temporal performance. This is what we explored next.

### α-β PAC indexes the precision in temporal performance

Cravo et al. (2011) reported θ-β PAC when participants maintained their internal state over a period of seconds to emit adaptive behavior. Building on this intuition, we hypothesized that PAC may reflect the ongoing anticipatory state during the endogenous elicitation of a duration goal. To test this, we studied how the observed PAC related to participants’ timing precision and accuracy. The α-β MI (8-12 Hz / 15-40 Hz) were averaged across sensors separately for each individual and for each experimental block, and entered as predictors in two regression models: the precision model (CVs) and the accuracy model (ERs).

In line with our predictions, we found that the strength of PAC significantly predicted CV [F(1, 65) = 8.6, *P* = 0.005; Fig. 4A]. The statistical analysis, based on Akaike Information Criterion (AIC, Wagenmakers, 2014), showed that the model containing α-β PAC as predictor was justified as compared to the model including only the intercept [ΔAIC = 6.3, *P* > 10^−15^]. Neither the inclusion of block factor, nor the inclusion of the interaction of α-β PAC with block factor in the model were warranted [ΔAIC = −7.3, *P* > 0.1; ΔAIC = −9.3, *P* > 0.1; respectively]). This indicated that in spite of variability in behavioral performance, the relationship between the precision as quantified by CV (Fig. 1C) and the α-β PAC was sustained throughout the entire experimental session.

**Fig. 4.**
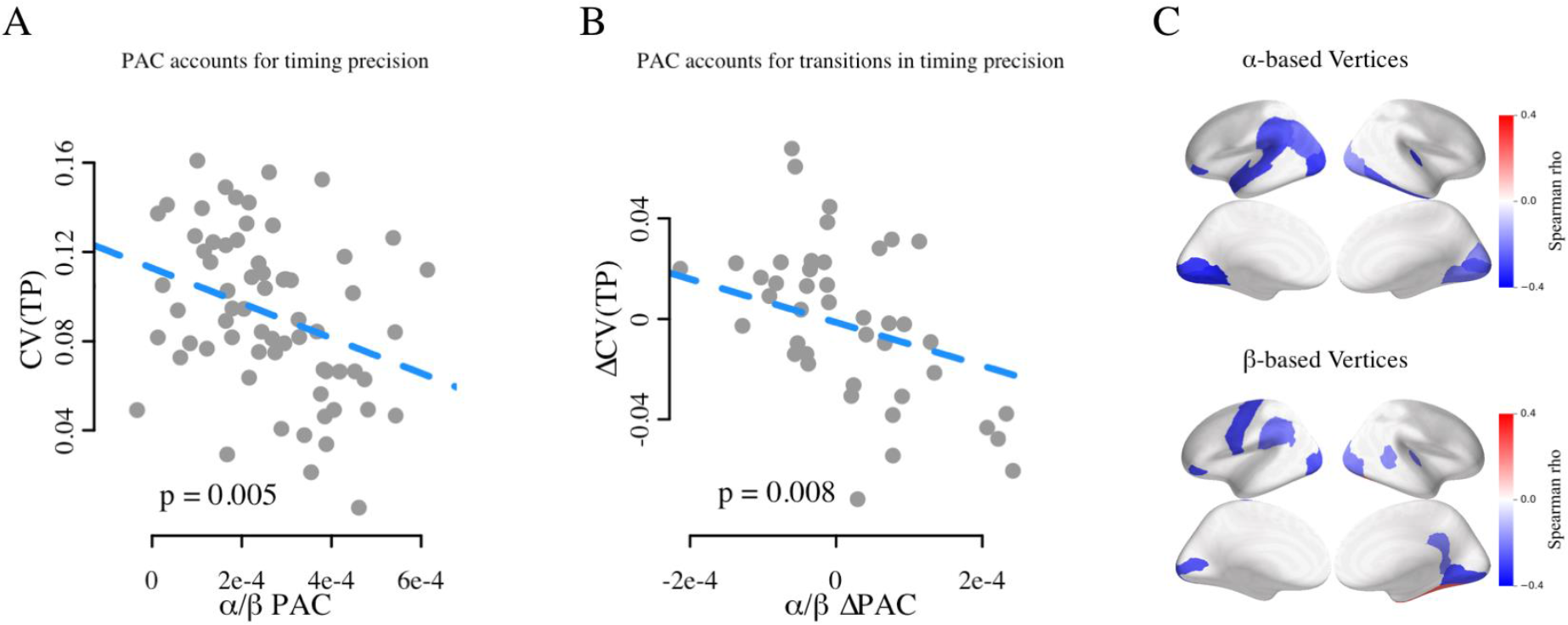
α-β PAC indexes the precision of temporal performance. (*A*) Over all blocks and participants the CV of temporal production was significantly correlated with the strength of *α* - *β* PAC, with no contribution of *α* or *α* power, suggesting that *α* - *β* PAC exclusively accounted for participants’ temporal precision. (B) The relative changes in CV and *α* - *β* PAC were correlated when participants switched from one block to another, suggesting that the transitions in *α* - *β* PAC strength between blocks were followed by transitions in CV. (C) Cortical source estimations of reported correlations between *α* - *β* PAC and CV.

To further test whether PAC indexed the precision of behavioral performance, we assessed whether transitions in α-β PAC between blocks (ΔPAC) could predict transitions in CV between blocks (ΔCV). For this, we subtracted the CV and the α-β MI between consecutive blocks (i.e., Block 2 – Block 1, Block 3 – Block 2, etc.). In line with the results of the first model, ΔPAC significantly predicted ΔCV [F(1, 53) = 7.7, *P* = 0.008; Fig. 4B], indicating that changes in precision (CV) between blocks could be accounted for by changes in PAC between blocks.

Previous studies on cross-frequency coupling have indicated that PAC estimation could be confounded during the estimation of phase and power frequencies (e.g. Aru et al., 2016). To thoroughly assess whether the predictive power of PAC with respect to CV was exclusive to oscillatory coupling, and not confounded by the power in α or β bands, we tested whether the inclusion of α power (Fig. S5A), β power (Fig. S5B), and α-β power ratio (Fig. S5C) was justified in the model predicting CV. We confirmed that the inclusion of *α*, *β*, and α-β ratio were not justified in the model predicting CV [Fig. ΔAIC = 0.1, *P* > 0.1; ΔAIC = 1.2, *P* > 0.05; ΔAIC = −1.9, *P* > 0.1; respectively].

Finally, we investigated the association of PAC with the precision of temporal behavior by correlating the values of the α-β MI obtained in each highlighted cortical region with the CVs: significant correlations between α-β MI and CV were found in the left motor areas, the left supramarginal gyrus, and occipito-parietal regions (Fig. 4C). Notably, the left supramarginal gyrus and pars orbitalis were both present in the α-based and in the β-based selection of vertices. The implication of these structures in the current timing task is consistent with previous reports (Coull, et al., 2004; Livesey et al., 2006, Wiener et al., 2010).

### α-β PAC and α power index temporal accuracy

α-β PAC was shown to correlate with the precision of temporal production (CV) across blocks with a stronger coupling indicating a smaller variance in time production. After investigating the *ad-hoc* hypothesis proposing that PAC could regulate the precision of information processing, we assessed whether α-β PAC related to the differences between produced and objective target intervals, i.e. to the accuracy. The accuracy assessed by ER indicated the distance of participant’s temporal production from an ideal observer’s performance (Fig. 1D). We found that PAC was significantly indicative of ER [F(1, 64) = 9.5, P = 0.003; Fig. 5A]. The AIC analysis showed that the model containing PAC as predictor was justified, as compared to the model including only the intercept [ΔAIC = 7.2, P = 0.002]. The inclusion of block factor to the model was also justified [ΔAIC = 7.4, P = 0.003] but no interactions between PAC and block factor were found [ΔAIC = 0.2, P > 0.1]. This result signified that changes in feedback did not affect the strength of PAC, and this was verified by directly contrasting blocks of 100% feedback with blocks of 15% feedback [F(1, 65) = 1.196, P = 0.2782]. It thus appeared that the relationship between the oscillatory coupling strength, as measured by PAC, and the accuracy of time estimation, as measured by ER, remained stable throughout the entire experimental session.

**Fig. 5.**
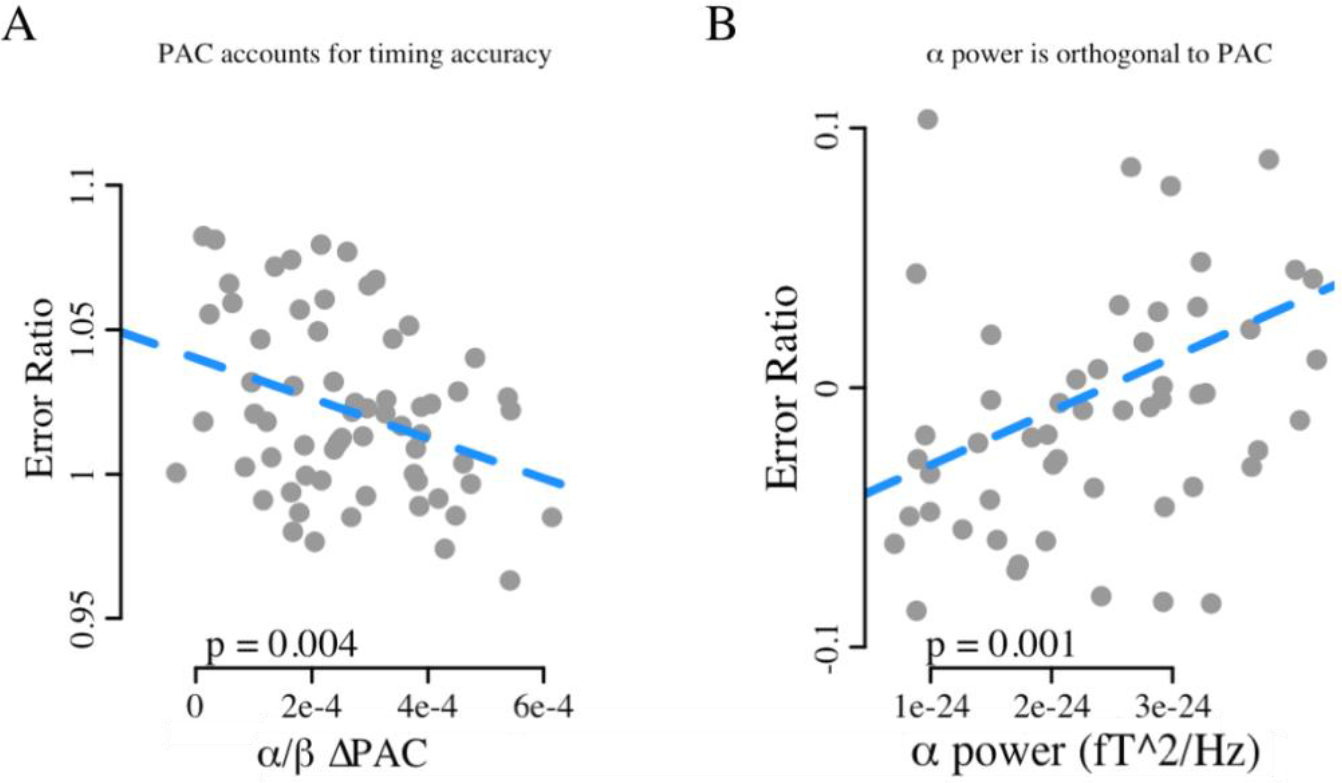
α-β PAC and α power index performance accuracy. (*A*) A decrease in ER) of time production was significantly correlated with a decrease in α-β coupling strength. (*B*) An increase in α power significantly correlated with an increase in ER. The scale in panel B differs from panel A because it reflects residuals in the model after accounting for α-β ΔPAC. These results indicate that accuracy is accounted for by α-β PAC and by a modulation of α power.

In the accuracy model, α power was also found to significantly contribute to the prediction of ER [t(57) = 3.7, *P* = 0.001] but the inclusion of PAC in the model remained significant [t(57) = −4.1, *P* < 0.001]. It is noteworthy that PAC and α power had opposite effects in predicting ER. Additionally, the inclusion of α power to the model was warranted by model comparison against the model including PAC [ΔAIC = 8.9, *P* < 0.001; Fig. 5B], whereas the inclusion of β (Fig. S5D) and α-β ratio (Fig. S5E) were not [ΔAIC = −10.9, *P* > 0.1; ΔAIC = −4.8, *P* > 0.1; respectively]. This observation confirmed the specificity of the contribution of PAC and α power to time estimation.

In line with the outcomes of a precision model, the accuracy model showed that a decrease in α-β PAC strength was linked to less accurate time productions. At first, the functional role of α-β PAC on accuracy seemed less clear than the one of precision; however, the decrease in accuracy could be caused by the loss of precision, with a higher chance of poorer time estimation.

### No monotonous association between PAC and time estimation

Although PAC has been proposed to mediate the integration of information across temporal scales for interval timing (Gu et al., 2015), the precise manner in which counting time would be implemented with oscillatory multiplexing remains difficult to elaborate (van Wassenhove, 2016). To clarify our results, we formulated a direct test of whether the generation of a time interval resulted from the online integration of endogenous information mediated by oscillatory multiplexing and investigated whether α-β PAC predicted timing behavior in an absolute manner between trials. In line with the common practice in the field (Kononowicz & Van Rijn, 2011), data were split in three bins as a function of participants’ performance: short, correct or long. The α-β MI was averaged across sensors for each individual and for each produced duration. A one-way repeated-measures ANOVA conducted with produced duration as factor (3 levels: short, correct, long) revealed no significant differences [F(2, 11)=0.657, *P* = 0.528; Fig. S6]. To investigate the likelihood of the null hypothesis, we ran the same analysis with Bayesian ANOVA: a Bayes Factor of 0.29 indicated that the data were 0.29 more likely to occur under the null hypothesis than under the alternative hypothesis. In other words, our results were 1/0.29 = 3.4 times more likely to occur under the null hypothesis, providing “substantial” (Jeffreys, 1961) or “moderate” (Lee & Wagenmakers, 2013) evidence that the α-β MI were independent of performance. We also tested whether the preferred phase of α-β coupling differed as a function of produced duration, but found no substantial evidence (Fig. S7). Our results thus provide no evidence for a specific role of cross-frequency coupling in duration estimation *per se*; rather, we find that α-β PAC indexes the precision with which the endogenous temporal goal is maintained across trials. One possible interpretation discussed below is that, from a neural network perspective, α may be regulating the β timing goal endogenously generated.

## Discussion

Using time-resolved neuroimaging, we investigated whether cross-frequency-coupling was implicated in the endogenous generation of time intervals. Our results provide strong evidence that α-β PAC leverages the precision of timing, but not the coding of a time interval. Specifically, we found that α-β PAC was indicative of the precision and accuracy of endogenous timing so that the stronger the α-β coupling, the more precise the performance (Fig. 6). These results suggest that α-β coupling indexes the precision with which information is endogenously maintained and transferred in the brain during a cognitive task.

**Fig. 6.**
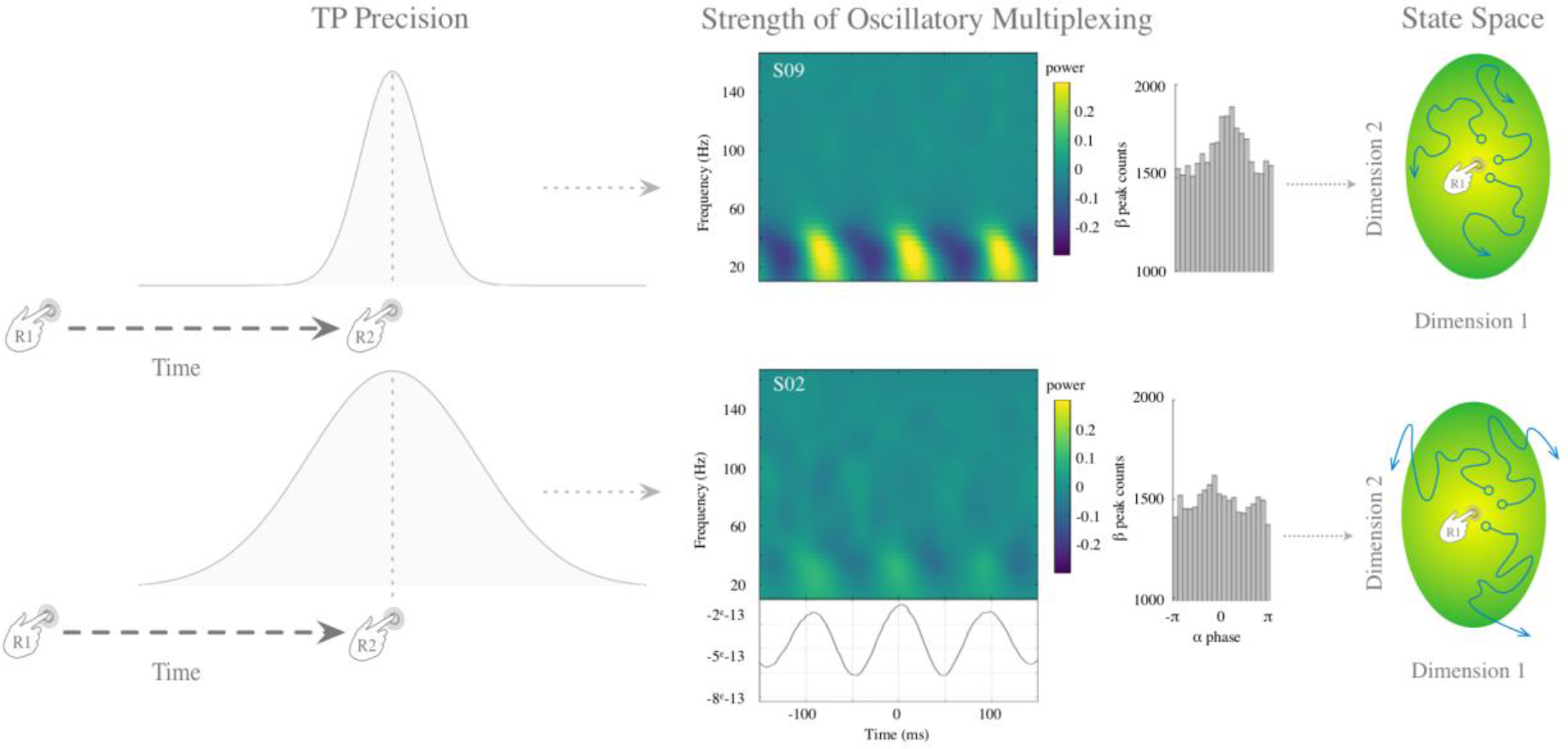
α-β PAC regulates the precision in time production. This schematic figure illustrates that the precision of temporal production in our study correlates with α-β PAC. Higher precision corresponds to a narrower nonuniform distribution of PAC (left top panel); lower precision to a broader non-uniform distribution (left bottom). Time-frequency plots for one individual with high precision (S09; top middle) and for one individual with low precision (S02; bottom middle) depicting the observed β power phase-locked to α phase: higher precision was associated with stronger α-β PAC, lower precision with weaker α-β PAC (middle top and bottom, respectively). The peak count distribution of β power maxima relative to the α phase are provided. The rightmost column depicts the working hypothesis that oscillatory coupling allows β oscillations to be reactivated, thereby endogenously modulating the timing goal encoded in the state space of synaptic configurations (see Discussion).

### α-β PAC is not related to motor preparation or learning

Due to fast progresses in uncovering the functions of cross-frequency coupling, accounting for the present findings with the existing evidence for the implication of PAC in motor preparation or learning would be tempting. However, several lines of evidence suggest that this is not the case. First, none of the frequency regimes described here conformed with previous reports: for instance, in intracranial recordings and ECoG, motor preparation was found to induce coupling in α/high-γ regimes, with an enhanced coupling during the preparation as compared to the movement execution phase (Combrison et al., 2017; Yanagisawa et al., 2012). Additionally, δ/α PAC contralateral to the side of the motor preparation was found to increase during movement *vs*. no movement (Kajihara et al. 2015). In situations of aberrant oscillatory regimes such as in patients with Parkinson, Hemptine et al. (2013) described an increased β/high γ PAC. Although we cannot be fully exhaustive, motor functions have typically been linked to different frequency coupling than the α-β PAC reported here. Another line of research reported the enhancement of PAC during learning (Tort et al., 2009). However, three pieces of evidence described in our study do not align well with the learning hypothesis, understood as progressive improvement in the measured neural or behavioral feature over the course of experiment (Kononowicz & Van Rijn, 2011; Tort et al., 2009). First, participants’ behavioral precision was stable over experimental blocks, indicating a stable level of performance in the course of the experiment. Second, the association between α-β PAC with behavioral precision was also stable over blocks, indicating no role for α-β PAC in potential learning-related effects. Third, under the learning hypothesis, it would be conceivable that different durations would entail different memory loads so that increased PAC would correlate with increased durations. We did not find such relationship between PAC and produced duration. Overall, our results cannot be directly interpreted in the context of learning effects linked to PAC (Tort et al., 2009) as this experimental design did not explicitly manipulate learning in time production.

### Maintenance of endogenous goal through α-β coupling

The specific role of α oscillations in cognition and information processing remains largely debated (Roux & Uhlhaas, 2013) but several observations show some consistencies. Among them, α oscillations have seminally been associated with the inhibition of irrelevant information (Klimesch et al., 2007; Jensen & Mazaheri, 2010; Haegens et al., 2012) and the maintenance of contents in working memory (Bonnefond & Jensen, 2012). In particular, the power of α oscillations has been shown to increase during cognitive tasks requiring internal engagement, which supports the protective role of α rhythms against exogenous distraction (Scheeringa et al. 2008). α oscillations have also been shown to play an active role in working memory maintenance (Palva et al., 2011) and in anticipation (Haegens et al., 2012; Praamstra et al., 2012). Both anticipation (Coull et al., 2013; Fortin, Bédard, & Champagne, 2005; Fortin & Massé, 2000) and maintenance of task goal (Lustig & Meck, 2001) are vital functions in time estimation. Hence, our observation of an active engagement of α oscillations in this task is consistent with prior literature.

β activity has been observed during time production and temporal expectations, suggesting that β oscillations may index a neural code for time estimation. Consistent with previous work using a motor timing task (Kononowicz and Van Rijn, 2015), increases in β power was found to index increases in time estimates. These findings have been extended to perceptual timing (Kulashekhar et al., 2016) and recently shown to be causally related to time estimation (Wiener et al., 2017). β oscillations have also been implicated in the generation of temporal predictions through sensorimotor interactions (Arnal et al., 2014; Fujioka et al., 2012) and involved in encoding of inter-tap duration (Bartolo et al., 2015). Hence, the active implication of β oscillations in this task is largely consistent with prior literature.

Altogether, our data are in line with a previous proposal suggesting that α and β regimes cooperate for content maintenance (Palva & Palva, 2007). This view support the recent ideas that α-driven PAC could provide a computational framework, in which α rhythms serve as a readout for relevant items (Roux & Uhlhaas, 2013; also see Nikulin, 2006). Specifically, the kind of time production paradigms used here relied on the anticipation of internally generated temporal codes and goals for duration production. In line with PAC shown during externally driven anticipatory states (Cohen et al., 2008; Cravo et al., 2011), our results show that enhanced PAC indexed more precise temporal production across blocks, and therefore appealed to the idea of coupling between neuronal regimes for the selection and the maintenance of neuronal representations (Palva & Palva, 2007). Given that the instantiation of PAC as α-β coupling found here was commensurate with the endogenous precision of a time goal, one hypothesis we could put forward is that β-based event computations may support endogenous temporal codes and goals.

### Specificity of α-β coupling to timing precision in the absence of external stimulation

A large body of work has suggested that PAC plays a functional role in maintaining information in working-memory and in attention. In working memory, PAC has been reported to operate in the θ-γ regimes (Axmacher et al., 2010; Fell & Axmacher, 2011; Alekseichuk et al., 2016), whereas modulations of attention may entail the α-γ regimes (Voytek et al., 2010; Roux & Uhlhaas, 2013). In particular, enhanced PAC has been associated with increased working memory load (Axmacher et al., 2012) and successful memory recollection in a cued task (Park et al., 2016). In both cases, γ activity was reported as the oscillatory component underlying information maintenance in working memory (Herrmann et al., 2004, but see Daume et al., 2017). Here, we found no compelling evidence for the *integration hypothesis* in duration estimation, which was rooted in these seminal findings.

Rather, our results were consistent with the *precision hypothesis*, which built on two studies showing a phase alignment of slow oscillations with the temporal task structure (Cravo et al., 2011; Samaha et al., 2015). For instance, Cravo et al. (2011) reported elevated PAC between two moments in time (temporal expectancy) for which it was perceptually beneficial for participants to maintain the same anticipation level. Both paradigms (Cravo et al., 2011; Samaha et al., 2015) relied on the individual’s anticipation to external sensory inputs, supporting the notion that anticipatory states can be maintained through α-β PAC. Our results extend these observations by showing that α-β PAC is relevant for temporal processing, not for the absolute time code in motor timing but for the precision it provides. More generally, it is also in line with a recent animal study suggesting that the precision of neural representations, including time intervals, is adjustable and may rely on multiple oscillators (Kheifets, et al., 2017).

It is also relevant that, with entrainment, the phase of low-frequency oscillations and beta activity has been reported to be relevant for timing. For example, δ-β PAC has been observed during the anticipation of visual sequences (Saleh et al., 2011) and related to the temporal resolution of perceptual systems (Arnal et al., 2014). High δ-β coupling has been associated with temporally accurate perception of an auditory stimulus embedded within a sequence (Arnal et al., 2014) and changes in the phase of δ oscillations has also been associated with explicit time order (Kösem et al., 2014). In all studies, unlike the current experimental approach, low-frequency oscillations were entrained by the sequence of sensory stimulations, and this may largely account for the differences between the δ-β reported before, and the α-β coupling observed here.

### α-β PAC as a signature of stabilization of synaptic state-space

α-β PAC has rarely been explored as a coupling configuration. Although the realm of possible coupling scenarios (e.g., Hyafil et al., 2015) preclude from drawing detailed conclusions on the computational architectures at the neurophysiological level (Canolty et al., 2010; Hyafil et al., 2015), our results highlight three features in broader neurophysiological terms. *First,* our results show that α and β generating networks are, to a certain extent, intertwined during a volitional task engaging no sensory stimulation and relying on self-generated timing (*i.e.* endogenous information). *Second*, that higher precision may arise from the maintenance of β-encoded information by the phase of α rhythm is in line with recent simulations using a 4-layers neuronal mass model (Sotero, 2015; 2016). This simulation reported the emergence of α-β PAC and, using information flow analysis, the phase of α oscillations was shown to stabilize β activity generated by fast spiking interneurons in layers 2/4 (Sotero, 2015; 2016). The reported directionality suggested that the phase of α oscillations may provide an optimal window for the reactivation of β-driven activity, consistent with the hypothesis that a global α network could regulate local β activity (e.g. Lee et al., 2013). Hence, one possible interpretation of our results is that β activity may reflect the neural state-space of synaptic configurations (Gavornik et al., 2009; Laje & Buonomano, 2013; Namboodiri et al., 2015) in cortical sources contributing to the maintenance of participants’ endogenous temporal goals (Fig. 6). β power, a marker of time estimation (Kononowicz and Van Rijn, 2015; Kulashekhar et al., 2016; Wiener et al., 2017), may also reflect the state-space of synaptic weights (Kononowicz & Penney, 2016), consistent with neural populations contributing to clocking mechanisms (Karmakar & Buonomano, 2004; Buonomano & Laje, 2010). This interpretation would be in line with recent suggestions that β synchronization provides a functional assembly-forming mechanism (Spitzer and Haegens, 2017; Lundqvist et al. 2016). Hence, what the α-β PAC as a marker of precision would signify in this context, is that in the absence of updating of the synaptic weights, the endogenous temporal goal may drift away from the optimal space as depicted in the trajectories distancing away from the colored area representing the optimal weights (Fig. 6). *Thirdly*, and importantly, the regulation of behavioral precision by α-β PAC need not be specific to timing. Here, the timing task we used enabled to move away from sensory-driven processes and uniquely focus on self-generated or endogenously driven activity. As such, the link between PAC and performance precision appears to capture the variance of computation in neuronal assemblies (Renart & Machens, 2014), and the precision with which assemblies act and represent task goals appear to translate into the temporal precision of behavioral performance.

### Conclusions

Oscillatory multiplexing, instantiated by phase-amplitude coupling, have been mainly studied in the context of working memory. Here, we showed the implication of a rarely studied coupling between α and β oscillations, in the self-generation of time intervals, and in the absence of sensory stimulation. The described α-β PAC supports the *precision* of temporal production. We suggest that α oscillations maintain and support the organization of β oscillations to keep track of endogenous temporal goals. In other words, our results provide strong evidence that α-β PAC leverages the precision of timing, but not absolute timing itself.

## Methods

### Participants

Nineteen right-handed volunteers (11 females, mean age: 24 years old) with no self-reported hearing/vision loss or neurological pathology took part in the experiment and received monetary compensation for participation. Each participant provided a written informed consent in accordance with the Declaration of Helsinki (2008) and the Ethics Committee on Human Research at Neurospin (Gif-sur-Yvette). The data of seven participants could not be included in the analysis due to the absence of anatomical MRI, problems with head positioning system, abnormal artifacts during MEG recordings, and two participants not finishing the experiment. These datasets were excluded *a priori* and were not visualized or inspected. Hence, the final sample comprised twelve participants (7 females, mean age: 24). All but two participants performed six experimental blocks; the first block was removed for one participant due to excessive artifacts, the last block was removed for another participant who did not conform to the requirements of the task.

### Stimuli and Procedure

Before the MEG acquisitions, participants were acquainted with the task by producing 1.45 s duration intervals, and reading written instructions explaining the full experimental procedures. A single trial consisted in producing a time interval followed by the self-estimation of the produced time interval. The feedback varied across blocks (block 1 and 4 included 100% feedback; blocks 2,3 and 5,6 included 15% feedback). Each trial started with the presentation of a “+” sign on the screen indicating to participants they could initiate an interval whenever they decided to (Fig. 1A). Participants initiated their production of the time interval with a brief button press (R1) once they felt relaxed and ready to start. Once they considered that a 1.45 s had elapsed, they terminated the interval by another brief button press (R2). To initiate and terminate their time production, participants were asked to press the top button on a Fiber Optic Response Pad (FORP, Science Plus Group, DE) using their right thumb. The “+” sign was removed from the screen during the estimation of the time interval to avoid any sensory cue or confounding responses in brain activity related to the production of the timed interval. Following the production of the time interval, participants were asked to self-estimate their time estimation using a continuous scale identical to the one used to provide feedback. The inter-trial interval between the end of the self-estimation and the first cursor display ranged between 1 s and 1.5 s.

Following the completion of the time interval, participants received feedback. The row of five symbols indicated the objective category of the time production tailored to each individual's time estimation. The feedback range was set to the value of the perceptual threshold estimated on a per individual basis during a task performed prior to MEG acquisition (mean population threshold = 0.223 s, SD = 0.111 s). A near correct time production yielded the middle ‘~’ symbol to turn green; a too short or too long time production turned the symbols ‘−’ or ‘+’ orange, respectively; a time production that exceeded these categories turned the symbols ‘− −’ or ‘++’ red. In Block 1, feedback was provided in all trials. In Block 2 and 3, feedback was randomly assigned to 15% of the trials. In Block 4, the target duration was increased to 1.45 + (*individual threshold*/2) and feedback was presented in all trials. On average, the new target duration was 1.56 s based on the average threshold. In Block 5 and 6, feedback was presented on 15% of trials. In Block 1 and 4, participants produced 100 trials; in Block 2, 3, 5, and 6, participants produced 118 trials.

Between all experimental blocks, participants were reminded to produce the 1.45 s target duration as accurately as possible and to maximize the number of correct trials in each block. During the estimation of individual thresholds, responses were collected using FORP, with the yellow button assigned to the index, the green button assigned to the middle finger and the red button assigned the ring finger of the right hand according to the position of the longest tone. Feedback was not provided to the participants.

### Simultaneous M/EEG recordings

The experiment was conducted in a dimly lit magnetically-shielded room located at Neurospin (CEA/DRF/Joliot) in Gif-sur-Yvette. Participants sat in an armchair with eyes opened looking at a projector screen. Electromagnetic brain activity was recorded using the whole-head Elekta Neuromag Vector View 306 MEG system (Neuromag Elekta LTD, Helsinki) equipped with 102 triple-sensors elements (two orthogonal planar gradiometers, and one magnetometer per sensor location) and the 64 native EEG system using Ag-AgCl electrodes (EasyCap, Germany) with impedances below 15 kΩ. Participants’ head position was measured before each block using four head-position coils placed over the frontal and the mastoid areas. The four head-position coils and three additional fiducial points (nasion, left and right pre-auricular areas) were digitized using a 3D digitizer (Polhemus, US/Canada) for subsequent co-registration of the individual’s anatomical MRI with brain recordings. MEG and EEG recordings were sampled at 1 kHz and band-pass filtered between 0.03 Hz and 330 Hz. The electro-occulograms (EOG, horizontal and vertical eye movements), -cardiograms (ECG), and -myograms (EMG) were simultaneously recorded. Feedback was presented using a PC running Psychtoolbox software (Brainard, 1997) in MATLAB (R2012, The Mathworks).

## Data Analysis

### M/EEG data preprocessing

MEG data were low-passed at 160 Hz, decimated at 333 Hz and epoched from −1.2s from the onset of the first button press (R1) up to 2 s after. Epochs were rejected if signal amplitudes exceeded 4 fT/cm for gradiometers and 5500 fT for magnetometers. Baseline correction was applied by subtracting the mean value ranging from −0.2 s to 0 s before R1. In this report, we focus on magnetometers and thus use the word “sensors” to refer to magnetometers.

### Power density spectrum (PSD) analysis

The PSD was computed using Welch’s method, between 1 and 45 Hz, with 800 ms length tapers, on a window from 0.4 to 1.2 s. PSD were averaged across all magnetometers, conditions and participants.

### Phase amplitude coupling calculation and statistical assessment

In order to prevent the contamination of the timed interval from evoked responses, we solely focused on the time segment from 0.4 s to 1.2 s following R1. PAC was assessed using the Modulation Index or MI (Tort et al., 2009), namely: raw data were band-pass filtered (slow-frequency bandwidth = 2 Hz, high-frequency bandwidth = 20 Hz); the instantaneous amplitude of the high-frequency and the phase of the slow-frequency were extracted from the Hilbert transform applied to the epoch data. To assess whether the distributions diverged from uniformity, the Kullback-Leibler (KL) distance was calculated then normalized to give the MI. The KL distance was estimated between histograms with 18 bins. The slow-frequency component ranged from 3.5 Hz to 14.1 Hz (step by 0.2 Hz) and the high-frequency range was from 14 Hz to 160 Hz (in step of 2 Hz). A comodulogram was computed for each sensor.

To assess the statistical significance of PAC at the individual level, the MI was compared to a surrogate distribution (n = 100) computed by shifting the low frequency signal by a minimum of 1 s, as has been previously proposed (Tort et al., 2010). A Z-score was calculated for each sensor and a Z-score higher than 4 (P = 3e−5) was reported as significant.

To assess whether PAC was specific to time production, we compared the MI computed during baseline from −0.8 s to 0 s prior to R1 to the MI computed during the produced interval ([0.4:1.2s]). Since participants were allowed to begin the task when ready after the cross display, we selected trials with at least 1.1 s between the cross onset and the first button press (R1) in order to avoid any contamination of visual evoked activity. On average 90 +/− 80 trials were retained. For 3 participants, the number of trials was not sufficient to compute a reliable MI (6, 15 and 21 trials; Fig. S6A). Nevertheless, the cluster-based permutation t-test was run on all individuals and similar results hold when the same analysis was carried out on 9 participants (Fig. S6B).

### M/EEG-aMRI coregistration

Anatomical Magnetic Resonance Imaging (aMRI) was used to provide high-resolution structural images of each individual’s brain. The anatomical MRI was recorded using a 3-T Siemens Trio MRI scanner. Parameters of the sequence were: voxel size: 1.0 × 1.0 × 1.1 mm; acquisition time: 466s; repetition time TR = 2300 ms; and echo time TE= 2.98 ms. Volumetric segmentation of participants’ anatomical MRI and cortical surface reconstruction was performed with the FreeSurfer software (http://surfer.nmr.mgh.harvard.edu/). A multi-echo FLASH pulse sequence with two flip angles (5 and 30 degrees) was also acquired (Fischl et al., 2004; Jovicich et al., 2006) to improve co-registration between EEG and aMRI. These procedures were used for group analysis with the MNE software (Gramfort et al., 2013; 2014). The co-registration of the M/EEG data with the individual’s structural MRI was carried out by realigning the digitized fiducial points with MRI slices. To insure reliable coregistration, an iterative refinement procedure was used to realign all digitized points with the individual’s scalp.

### MEG source reconstruction for PAC analysis

Individual forward solutions for all source locations located on the cortical sheet were computed using a 3-layers boundary element model (BEM) constrained by the individual’s aMRI. Cortical surfaces extracted with FreeSurfer were sub-sampled to 10,242 equally spaced sources on each hemisphere (3.1 mm between sources). The noise covariance matrix for each individual was estimated from the baseline activity of all trials and all conditions. The forward solution, the noise covariance and source covariance matrices were used to calculate the dSPM estimates (Dale et al., 2000). The inverse computation was done using a loose orientation constraint (loose = 0.4, depth = 0.8) on the radial component of the signal. Individuals’ current source estimates were registered on the Freesurfer average brain for surface based analysis and visualization. Once the time-resolved signals were reconstructed in cortical space, PAC was computed for each cortical label (‘aparc’ parcellation). As aparc labels can be quite large, we selected within each label with the largest values of α or β power. The label time courses were than treated as single trials.

### Experimentally-driven conditions and correlation analyses

Analyses were performed on the basis of the objective performance in time production classified as short, correct, or long separately for each experimental block. Computing these three conditions within a block focused the analysis on local variations of brain activity as a function of objective performance. Epochs were concatenated across all 6 blocks for the analyses based on time production performance. The number of trials was equalized between short, correct and long conditions, leading to 168 trials (SD = 58) per condition. Additionally, the correlational analyses investigating precision and accuracy of timing processes has been also extended to per block basis to gain better resolution in the fluctuation of these component of behavior over time. Precision has been computed as coefficient of variation over TP on a per block and participant basis, whereas accuracy has been calculated as by subtracting mean TP from the target duration in a given block. Note that in the first three blocks target duration was 1.45, however it has been dynamically assigned in the blocks from four to six.

Sensor level analyses were performed using linear regression and model comparison framework to check if other factor other than PAC were needed to explain behavior. All statistical analyses have been performed in R v3.3.2 statistical programming language (R Development Core 2008). For illustrative purposes, correlations in the source space were computed using Spearman correlations. Byesian ANOVA (Rouder et al., 2012) was performed using *BayesFactor* R package.

### Calculating precision and accuracy metrics and accounting for inflated degrees of freedom

Precision was quantified by the Coefficient of Variation (CV): the CV were calculated by dividing the standard deviation by the mean duration production. Accuracy was quantified by Error Ratio (ER): ER was calculated by dividing the mean temporal production in a given set of data by the target interval in that set. Both metrics were computed separately per block and per participant. As splitting the data per block and per individual could inflate the degrees of freedom, we accounted for this in our analysis by assessing the impact of block factor in the overall model fit by model comparison approach. In none of the analyses, the block factor altered the main effects and this is systematically and explicitly reported in our Results section when relevant.

### Data standardizing for regression model

As regression tests require a Gaussian distribution of the data, wherever applicable, and on the basis of Shapiro-Wilk normality test, the data were transformed using the Lambert W function. The Lambert W function provides an explicit inverse transformation, which removes heavy tails from the observed data (Georg, 2011, 2015). First, the data were transformed into a heavy tailed form using log-likelihood decomposition. Subsequently the heavy tailed form was back transformed into a Gaussian distribution. All transformations were performed using *LambertW* R package.

## Supplementary Captions and Figures

**Fig. S1.**
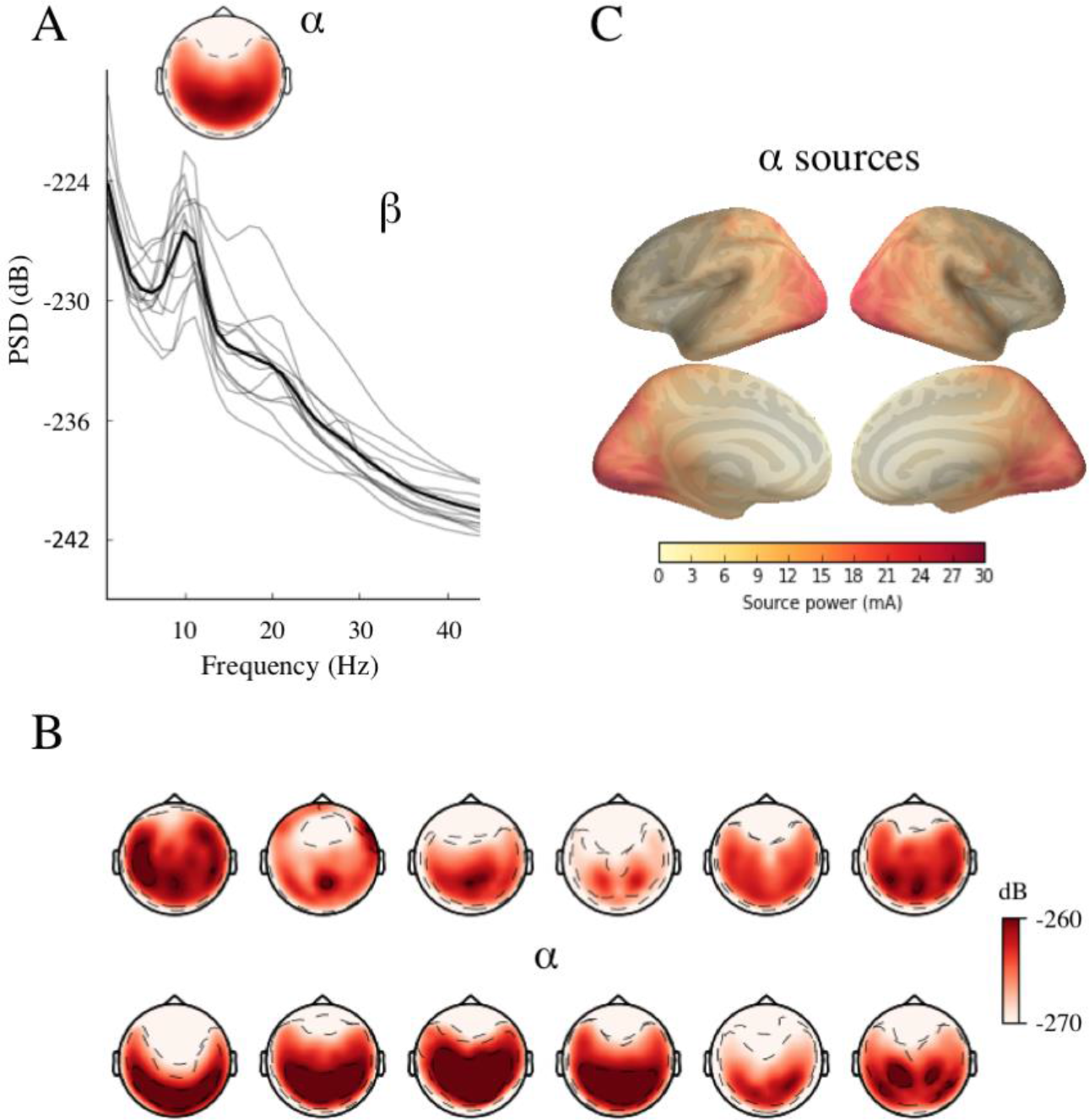
Power during time production. (A) The power spectrum density (PSD) was computed during the produced interval (0.4 - 1.2s) and averaged across channels, conditions and participants. The average PSD (thick line) across individuals (grey lines) show a clear α peak around 10 Hz. The topomap shows the PSD averaged between 8 and 12 Hz. (B) The individual topographic maps for α PSD (C) and grand-average source reconstruction reveal an occipito-parietal distribution of α power.

**Fig. S2.**
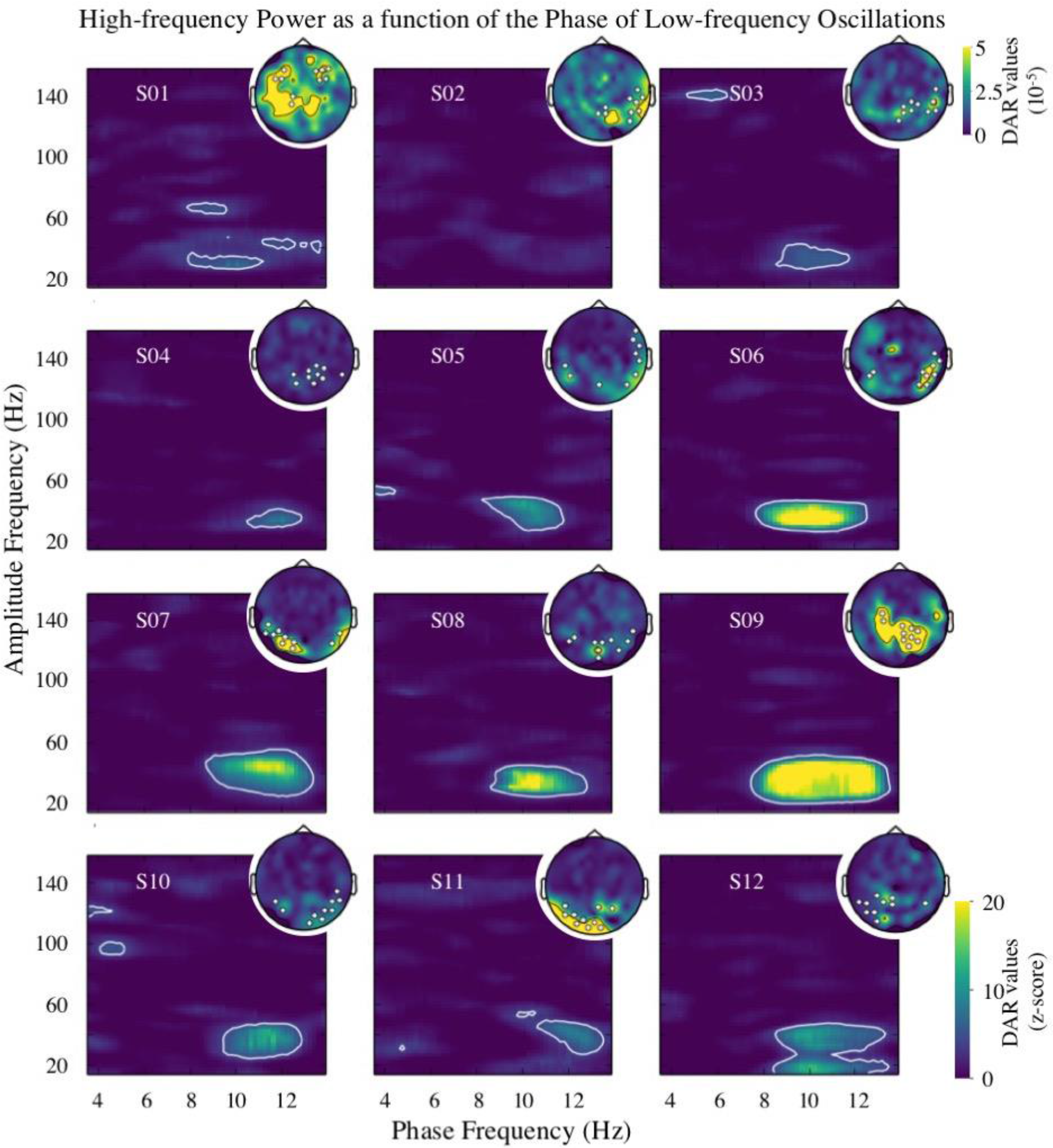
α-β PAC during time production computed with Driven Auto-Regressive model. We replicated the individual results on α-β PAC (Fig. 2A) with a novel method based on driven auto-regressive (DAR) model. Full comodulogram of Z-scored DAR values are plotted for each individual and topographic maps of α-β DAR values are plotted in the right insets. The white contour corresponds to Z-score values above 4. For DAR analysis, we kept the same set of sensors as in Fig 2A. In participant S02, sensors showing maximal PAC with Tort’s method (highlighted in white) did not match the sensors showing maximal PAC with DAR models; this spatial discrepancy explains why no significant α-β PAC was observed for S02.

**Fig. S3.**
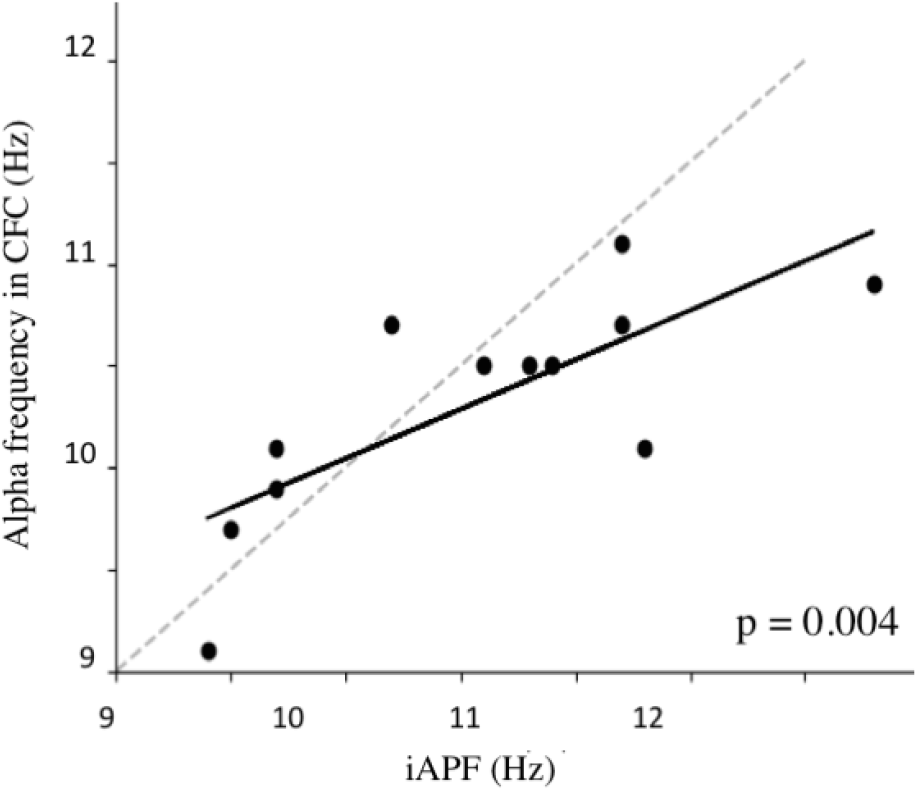
Individual α peak frequency correlates with α frequency in α-β PAC. To ensure that the α rhythm captured in the PSD was involved in PAC computation, we correlated the individual α peak frequency with the frequency corresponding to a maximal α-β MI [R = 0.763, *P* = 0.004].

**Fig. S4.**
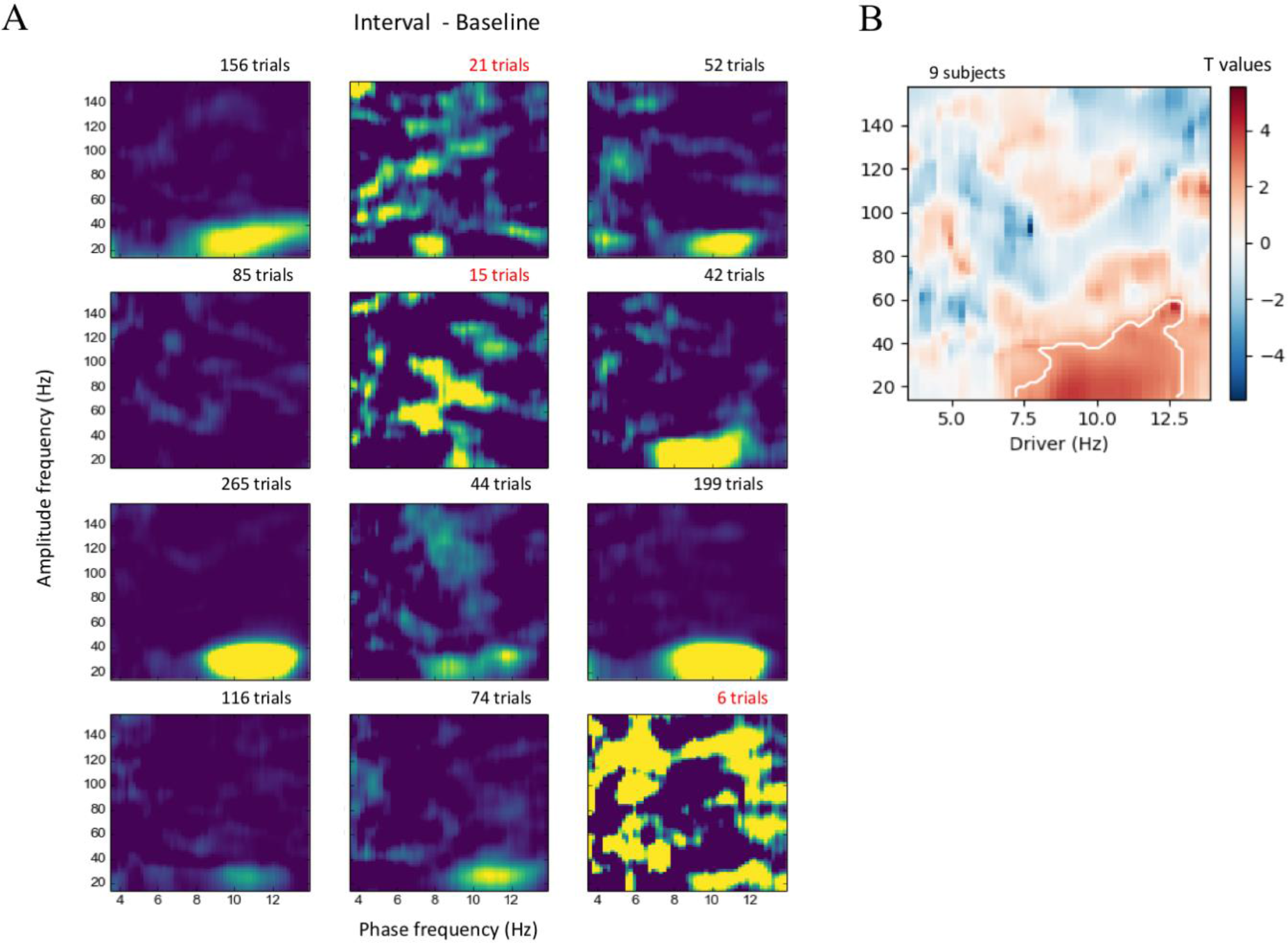
α-β PAC comparison against volitional motor preparation. (A) MI for each individual, computed on the trials that had available 0.8 s baseline interval. The number of available trials is provided above each comodulogram. The average topomap of α-β MI is plotted in the right inset. Participants with too few trials were both considered and disregarded for the analysis and did not affect the results (see text). (B) α-β PAC compared against baseline for 9 participants with a sufficient number of trials. As for the whole set of 12 subjects, α-β coupling was also significantly higher during the produced time interval as compared to the volitional motor preparation.

**Fig. S5.**
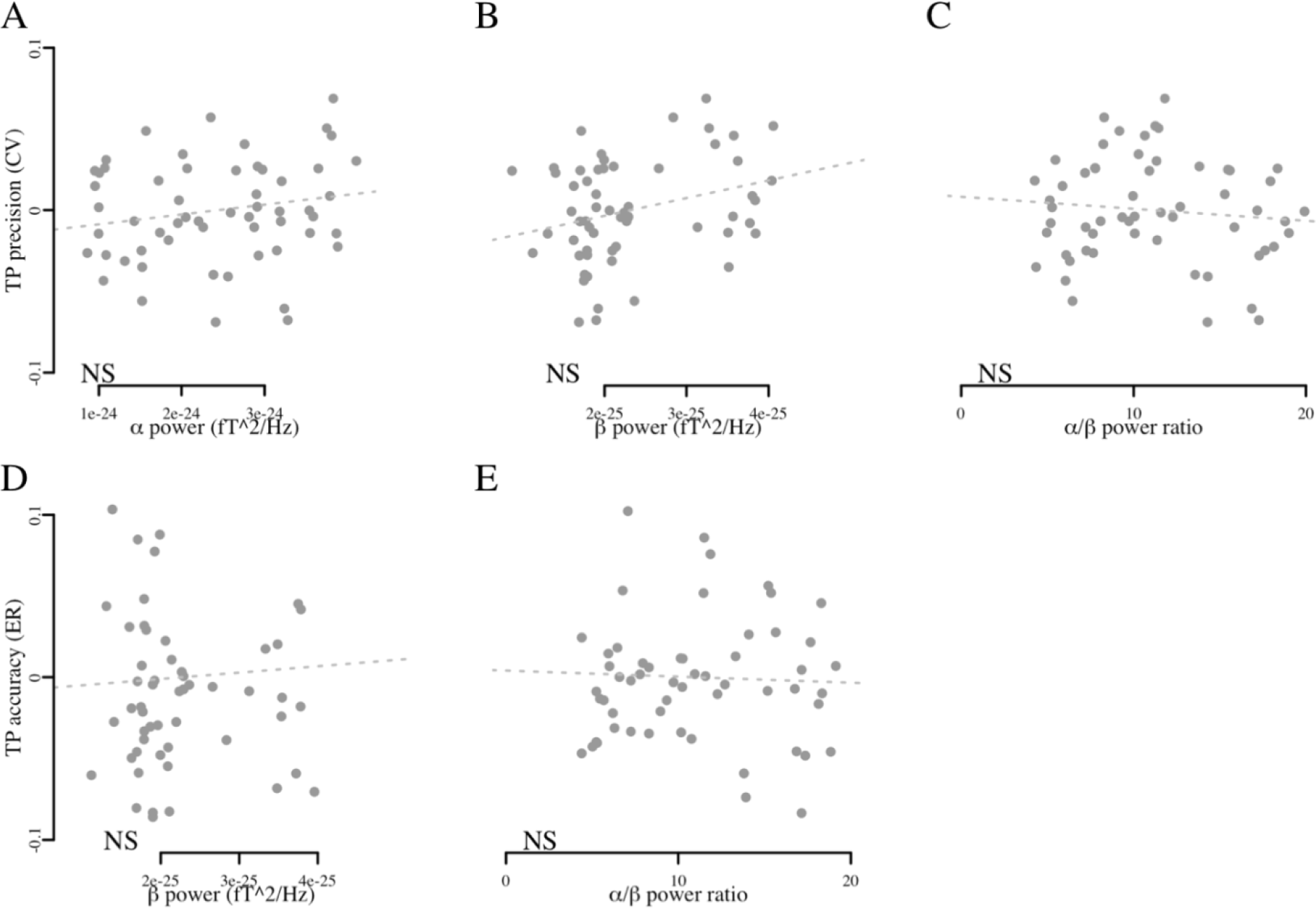
α and β power do not contribute to temporal performance precision. (A,B,C) Over all blocks and participants, the CVs of time interval did not correlate with the strength of α and β power or their ratio. In the absence of significant contributions of α (D) or β (E) power to ER in time production (TP), TP did not correlate with the strength of β power and α-β ratio.

**Fig. S6.**
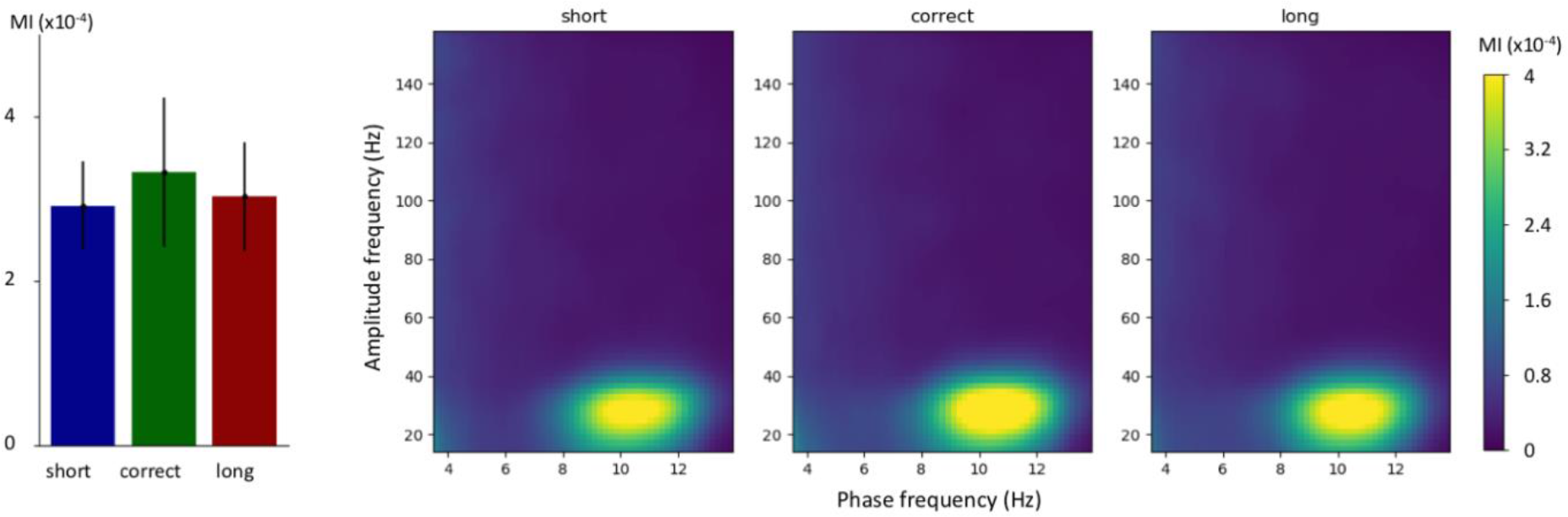
α-β coupling strength does not index absolute time production. Trials were split in three conditions according to the length of the temporal production (short in blue, correct in green, long in red). Error bars are 2 s.e.m.

**Fig. S7.**
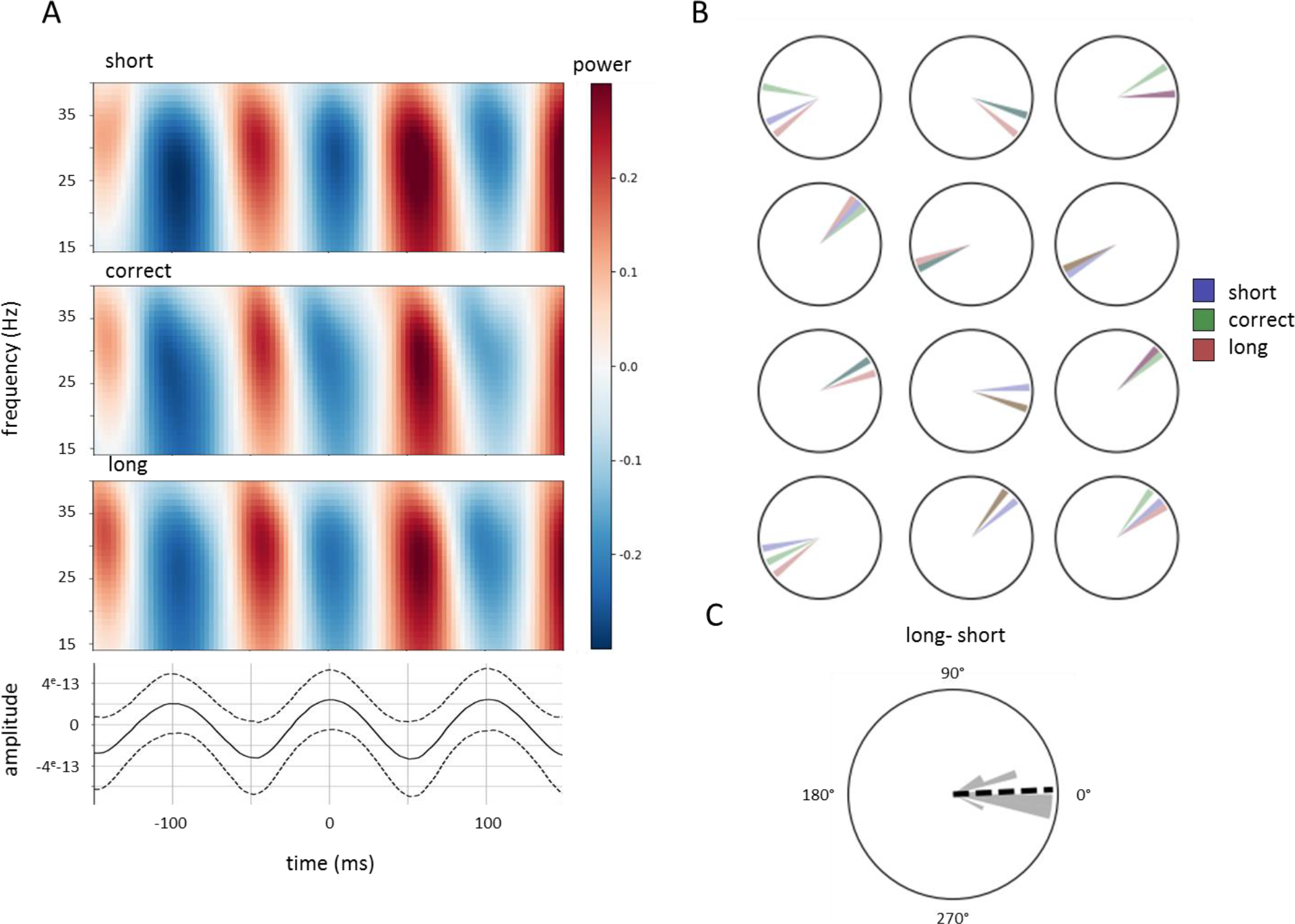
The phase relationship between α and β does not predict self-estimated time production. **A.** The time series were locked on the peak of the α oscillations (bottom, dark line, dashed line is std) and beta power (15-40Hz) were computed for each self-estimation duration. For illustration, the time-frequency phase-locked to the α peak is shown for the MEG sensor showing maximal α-β MI for one representative participant (S06). The increase in β occurred at the ascending slope of the α through for 4 participants, as shown here for one of them but at the descending slope for 8 others. **B.** The α phase at which β power was maximal was extracted for each participant (circular histogram) and each duration category (blue: short, green: correct, red: long). **C.** The phase difference between long and short condition was plotted for each participant. The length of the bar represents the number of participants with the same phase value. A Rayleigh test indicated that the phase difference between long and short categorized was not uniform (p < 10-3, mean = 2.7°) but the mean difference between long and short self-estimated duration did not significantly differ (t = 0.574, p = 0.578).

**Table S1.** Individual α peak frequency extracted from the PSD and individual α-β PAC corresponding to their maximal MI. CFC = cross-frequency coupling (here, equivalent to PAC).

| Participants | Alpha power peak (Hz) | Alpha CFC peak (Hz) | Beta CFC peak (Hz) |
| --- | --- | --- | --- |
| S01 | 9.40 | 9.1 | 18 |
| S02 | 11.20 | 11.1 | 32 |
| S03 | 9.70 | 9.9 | 28 |
| S04 | 11.30 | 10.1 | 24 |
| S05 | 9.50 | 9.7 | 26 |
| S06 | 9.70 | 10.1 | 28 |
| S07 | 12.30 | 10.9 | 30 |
| S08 | 10.60 | 10.5 | 30 |
| S09 | 10.90 | 10.5 | 32 |
| S10 | 10.20 | 10.7 | 28 |
| S11 | 11.20 | 10.7 | 26 |
| S12 | 10.80 | 10.5 | 24 |

## Acknowledgments

This work was supported by the Paris-Saclay IDEX NoTime to V.D., A.G. and V.vW, the ERC Starting Grant MINDTIME ERC-YStG-263584 grant to V.vW, the ERC Starting Grant SLAB ERC-YStG-676943 to A.G., and the ANR-16-CE37-0004-04 AutoTime to V.D. and V.vW. The funders had no role in study design, data collection and analysis, decision to publish, or preparation of the manuscript. Preliminary results of this work were presented at the *International Cognitive Neuroscience* workshop (Amsterdam, Netherlands, 2017).

## References

Akam, T., & Kullmann, D. M. (2014). Oscillatory multiplexing of population codes for selective communication in the mammalian brain. Nature Reviews. Neuroscience, 15(2), 111.

Arnal, L. H., Doelling, K. B., & Poeppel, D. (2014). Delta--beta coupled oscillations underlie temporal prediction accuracy. Cerebral Cortex, bhu103.

Axmacher, N., Henseler, M. M., Jensen, O., Weinreich, I., Elger, C. E., & Fell, J. (2010). Cross-frequency coupling supports multi-item working memory in the human hippocampus. Proceedings of the National Academy of Sciences, 107(7), 3228–3233.

Bartolo, R., & Merchant, H. (2015). B oscillations are linked to the initiation of sensory-cued movement sequences and the internal guidance of regular tapping in the monkey. The Journal of Neuroscience, 35(11), 4635–4640.

Bonnefond, M., & Jensen, O. (2012). Alpha oscillations serve to protect working memory maintenance against anticipated distracters. Current Biology, 22(20), 1969–1974.

Brainard, D. H. (1997) The Psychophysics Toolbox, Spatial Vision 10, 433–436.

Buonomano, D. V., & Laje, R. (2010). Population clocks: motor timing with neural dynamics. Trends in cognitive sciences, 14(12), 520–527.

Buzsáki, G., & Draguhn, A. (2004). Neuronal oscillations in cortical networks. Science, 304(5679), 1926–1929.

Buzsáki, G. (2010). Neural syntax: Cell assemblies, synapsembles, and readers. Neuron, 68(3), 362–385.

Canolty, R. T., Edwards, E., Dalal, S. S., Soltani, M., Nagarajan, S. S., Kirsch, H. E., … Knight, R. T. (2006). High gamma power is phase-locked to theta oscillations in human neocortex. Science, 313(5793), 1626–1628.

Canolty, R. T., & Knight, R. T. (2010). The functional role of cross-frequency coupling. Trends in Cognitive Sciences, 14(11), 506–515.

Chakravarthi, R., & VanRullen, R. (2012). Conscious updating is a rhythmic process. Proceedings of the National Academy of Sciences, 109(26), 10599–10604.

Cohen, M. X., Axmacher, N., Lenartz, D., Elger, C. E., Sturm, V., & Schlaepfer, T. E. (2009). Good vibrations: Cross-frequency coupling in the human nucleus accumbens during reward processing. Journal of Cognitive Neuroscience, 21(5), 875–889.

Combrisson, E., Perrone-Bertolotti, M., Soto, J. L., Alamian, G., Kahane, P., Lachaux, J. -P., … Jerbi, K. (2017). From intentions to actions: Neural oscillations encode motor processes through phase, amplitude and phase-amplitude coupling. NeuroImage, 147, 473–487.

Coull, J. T., Davranche, K., Nazarian, B., & Vidal, F. (2013). Functional anatomy of timing differs for production versus prediction of time intervals. Neuropsychologia, 51(2), 309–319.

Coull, J. T., Vidal, F., Nazarian, B., & Macar, F. (2004). Functional anatomy of the attentional modulation of time estimation. Science, 303(5663), 1506–8.

Cravo, A. M., Rohenkohl, G., Wyart, V., & Nobre, A. C. (2011). Endogenous modulation of low frequency oscillations by temporal expectations. Journal of Neurophysiology, 106(6), 2964–2972.

Dale, A. M., Liu, A. K., Fischl, B. R., Buckner, R. L., Belliveau, J. W., Lewine, J. D., et al. (2000). Dynamic statistical parametric mapping: Combining fMRI and MEG for high-resolution imaging of cortical activity. Neuron, 26, 55 – 67.

Goerg, G. M. (2011). Lambert W random variablesa new family of generalized skewed distributions with applications to risk estimation. The Annals of Applied Statistics, 2197–2230.

Goerg, G. M. (2015). The lambert way to gaussianize heavy-tailed data with the inverse of tukeysh transformation as a special case. The Scientific World Journal, 2015.

Gramfort, A., Luessi, M., Larson, E., Engemann, D., Strohmeier, D., Brodbeck, C., Goj, R., Jas, M., Brooks, T., Parkkonen, L., Hämäläinen, M., MEG and EEG data analysis with MNE-Python, Frontiers in Neuroscience, Volume 7, 2013, ISSN 1662-453X

Gramfort, A., Luessi, M., Larson, E., Engemann, D., Strohmeier, D., Brodbeck, C., Parkkonen, L., Hämäläinen, M., MNE software for processing MEG and EEG data, NeuroImage, Volume 86, 1 February 2014, Pages 446–460, ISSN 1053-8119

Daume, J, Gruber, T, Engel, A. K., Friese U. (2017) Phase-amplitude coupling and long-range phase synchronization reveal frontotemporal interactions during visual working memory. Journal of Neuroscience, 37(2), 313–322.

De Hemptinne, C., Swann, N. C., Ostrem, J. L., Ryapolova-Webb, E. S., San Luciano, M., Galifianakis, N. B., & Starr, P. A. (2015). Therapeutic deep brain stimulation reduces cortical phase-amplitude coupling in parkinson's disease. Nature Neuroscience, 18(5), 779–786.

Fischl, B., Salat, D. H., van der Kouwe, A. J., Makris, N., Ségonne, F., Quinn, B. T., & Dale, A. M. (2004). Sequence-independent segmentation of magnetic resonance images. Neuroimage, 23, 69–84.

Fortin, C., & Massé, N. (2000). Expecting a break in time estimation: Attentional timesharing without concurrent processing. Journal of Experimental Psychology: Human Perception and Performance, 26(6), 1788.

Fortin, C., Bédard, M. C., & Champagne, J. (2005). Timing during interruptions in timing. J Exp Psychol Hum Percept Perform, 31(2), 276–88.

Fries, P. (2015). Rhythms for cognition: Communication through coherence. Neuron, 88(1), 220–235.

Gavornik, J. P., Shuler, M. G. H., Loewenstein, Y., Bear, M. F., & Shouval, H. Z. (2009). Learning reward timing in cortex through reward dependent expression of synaptic plasticity. Proceedings of the National Academy of Sciences, 106(16), 6826–6831.

Giraud, A. -L., & Poeppel, D. (2012). Cortical oscillations and speech processing: Emerging computational principles and operations. Nature Neuroscience, 15(4), 511–517.

Gu, B. -M., van Rijn, H., & Meck, W. H. (2015). Oscillatory multiplexing of neural population codes for interval timing and working memory. Neuroscience & Biobehavioral Reviews, 48, 160–185.

Hyafil, A., Giraud, A. -L., Fontolan, L., & Gutkin, B. (2015). Neural cross-frequency coupling: Connecting architectures, mechanisms, and functions. Trends in Neurosciences, 38(11), 725–740.

Jacobs, J., Kahana, M. J., Ekstrom, A. D., & Fried, I. (2007). Brain oscillations control timing of single-neuron activity in humans. The Journal of Neuroscience, 27(14), 3839–3844.

Jeffreys, H. (1961). Theory of probability (3rd ed.). Oxford: Oxford University Press, Clarendon Press.

Jensen, O., & Colgin, L. L. (2007). Cross-frequency coupling between neuronal oscillations. Trends in Cognitive Sciences, 11(7), 267–269.

Jensen, O., & Mazaheri, A. (2010). Shaping functional architecture by oscillatory alpha activity: Gating by inhibition. Frontiers in Human Neuroscience, 4.

Jensen, O., Gips, B., Bergmann, T. O., & Bonnefond, M. (2014). Temporal coding organized by coupled alpha and gamma oscillations prioritize visual processing. Trends in Neurosciences, 37(7), 357–369.

Jones, S. R. (2016). When brain rhythms aren't rhythmic: Implication for their mechanisms and meaning. Current Opinion in Neurobiology, 40, 72–80.

Jovicich, J., Czanner, S., Greve, D., Haley, E., van der Kouwe, A., Gollub, R., … MacFall, J. (2006). Reliability in multi-site structural MRI studies: Effects of gradient non-linearity correction on phantom and human data. Neuroimage, 30(2), 436–443.

Karmarkar, U. R., & Buonomano, D. V. (2007). Timing in the absence of clocks: encoding time in neural network states. Neuron, 53(3), 427–438.

Kawai, R., Markman, T., Poddar, R., Ko, R., Fantana, A. L., Dhawale, A. K., … Ölveczky, B. P. (2015). Motor cortex is required for learning but not for executing a motor skill. Neuron, 86(3), 800–812.

Kheifets, A., Freestone, D., & Gallistel, C. R. (2017). Theoretical implications of quantitative properties of interval timing and probability estimation in mouse and rat. Journal of the Experimental Analysis of Behavior, 108(1), 39–72.

Klimesch, W., Sauseng, P., & Hanslmayr, S. (2007). EEG alpha oscillations: The inhibition--timing hypothesis. Brain Research Reviews, 53(1), 63–88.

Kononowicz, T. W., & Van Rijn, H. (2011). Slow potentials in time estimation: The role of temporal accumulation and habituation. Frontiers in Integrative Neuroscience, 5(48).

Kononowicz, T. W., & van Rijn, H. (2015). Single trial beta oscillations index time estimation. Neuropsychologia, 75, 381–389.

Kononowicz, T. W., & van Wassenhove, V. (2016). In search of oscillatory traces of the internal clock. Frontiers in Psychology, 7.

Kononowicz, T. W., & Penney, T. B. (2016). The contingent negative variation (CNV): Timing isnt everything. Current Opinion in Behavioral Sciences, 8, 231–237.

Kösem, Anne, Alexandre Gramfort, and Virginie van Wassenhove. "Encoding of event timing in the phase of neural oscillations." NeuroImage 92 (2014): 274–284.

Kulashekhar, S., Pekkola, J., Palva, J. M., & Palva, S. (2016). The role of cortical beta oscillations in time estimation. Human Brain Mapping, 37(9), 3262–3281.

Laje, R., & Buonomano, D. V. (2013). Robust timing and motor patterns by taming chaos in recurrent neural networks. Nature neuroscience, 16(7), 925–933.

Dupré La Tour, T. D., Tallot, L., Grabot, L., Doyere, V., van Wassenhove, V., Grenier, Y., & Gramfort, A. (2017). Non-linear Auto-Regressive Models for Cross-Frequency Coupling in Neural Time Series. bioRxiv, 159731.

Lee, J. H., Whittington, M. A., & Kopell, N. J. (2013). Top-down beta rhythms support selective attention via interlaminar interaction: a model. PLoS computational biology, 9(8), e1003164.

Lee, M. D., & Wagenmakers, E. -J. (2014). Bayesian cognitive modeling: A practical course. Cambridge University Press.

Lisman, J. E., & Jensen, O. (2013). The theta-gamma neural code. Neuron, 77(6), 1002–1016.

Livesey, A. C., Wall, M. B., & Smith, A. T. (2007). Time perception: Manipulation of task difficulty dissociates clock functions from other cognitive demands. Neuropsychologia, 45(2), 321–331.

Lundqvist, M., Rose, J., Herman, P., Brincat, S. L., Buschman, T. J., & Miller, E. K. (2016). Gamma and beta bursts underlie working memory. Neuron, 90(1), 152–164.

Lustig, C., & Meck, W. H. (2001). Paying attention to time as one gets older. Psychological Science, 12(6), 478–484.

MATLAB and Statistics Toolbox Release 2012b, The MathWorks, Inc., Natick, Massachusetts, United States.

Namboodiri, V. M. K., Huertas, M. A., Monk, K. J., Shouval, H. Z., & Shuler, M. G. H. (2015). Visually cued action timing in the primary visual cortex. Neuron, 86(1), 319–330.

Nikulin, V. V., & Brismar, T. (2006). Phase synchronization between alpha and beta oscillations in the human electroencephalogram. Neuroscience, 137(2), 647–657.

Palva, S., & Palva, J. M. (2007). New vistas for α-frequency band oscillations. Trends in Neurosciences, 30(4), 150–158.

R Core Team (2012). R: A language and environment for statistical computing. R Foundation for Statistical Computing, Vienna, Austria. ISBN 3-900051-07-0, URL http://www.R-project.org/

Renart, A., & Machens, C. K. (2014). Variability in neural activity and behavior. Current Opinion in Neurobiology, 25, 211–220.

Rouder, J. N., Morey, R. D., Speckman, P. L., & Province, J. M. (2012). Default bayes factors for ANOVA designs. Journal of Mathematical Psychology, 56(5), 356–374.

Samaha, J., Bauer, P., Cimaroli, S., & Postle, B. R. (2015). Top-down control of the phase of alpha-band oscillations as a mechanism for temporal prediction. Proceedings of the National Academy of Sciences, 112(27), 8439–8444.

Scheeringa, R., Petersson, K. M., Oostenveld, R., Norris, D. G., Hagoort, P., & Bastiaansen, M. C. (2009). Trial-by-trial coupling between EEG and BOLD identifies networks related to alpha and theta EEG power increases during working memory maintenance. Neuroimage, 44(3), 1224–1238.

Sotero, R. C. (2016). Topology, cross-frequency, and same-frequency band interactions shape the generation of phase-amplitude coupling in a neural mass model of a cortical column. PLoS Computational Biology, 12(11), e1005180.

Sotero, R. C. (2015). Modeling the generation of phase-amplitude coupling in cortical circuits: From detailed networks to neural mass models. BioMed Research International, 2015.

Spitzer, B., & Haegens, S. (2017). Beyond the status quo: A role for beta oscillations in endogenous content (re-) activation. ENeuro, ENEURO-0170.

Tort, A. B., Komorowski, R. W., Manns, J. R., Kopell, N. J., & Eichenbaum, H. (2009). Theta--gamma coupling increases during the learning of item--context associations. Proceedings of the National Academy of Sciences, 106(49), 20942–20947.

Tort, A. B., Kramer, M. A., Thorn, C., Gibson, D. J., Kubota, Y., Graybiel, A. M., & Kopell, N. J. (2008). Dynamic cross-frequency couplings of local field potential oscillations in rat striatum and hippocampus during performance of a t-maze task. Proceedings of the National Academy of Sciences, 105(51), 20517–20522.

Tort, A. B., Komorowski, R., Eichenbaum, H., & Kopell, N. (2010). Measuring phase-amplitude coupling between neuronal oscillations of different frequencies. Journal of Neurophysiology, 104(2), 1195–1210.

van Wassenhove, V. (2016). Temporal cognition and neural oscillations. Current Opinion in Behavioral Sciences, 8, 124–130.

Voytek, B., Canolty, R. T., Shestyuk, A., Crone, N. E., Parvizi, J., & Knight, R. T. (2010). Shifts in gamma phase--amplitude coupling frequency from theta to alpha over posterior cortex during visual tasks. Frontiers in Human Neuroscience, 4.

Wagenmakers, E. -J., & Farrell, S. (2004). AIC model selection using akaike weights. Psychonomic Bulletin & Review, 11(1), 192–196.

Yanagisawa, T., Yamashita, O., Hirata, M., Kishima, H., Saitoh, Y., Goto, T., … Kamitani, Y. (2012). Regulation of motor representation by phase--amplitude coupling in the sensorimotor cortex. The Journal of Neuroscience, 32(44), 15467–15475.

